# Femora Nutrient Foramina and Aerobic Capacity in Giant Extinct Xenarthrans

**DOI:** 10.1101/2023.09.27.559456

**Authors:** Luciano Varela, P. Sebastián Tambusso, Richard A. Fariña

**Affiliations:** Departamento de Paleontología, Facultad de Ciencias, Universidad de la República, Iguá 4225, 11400, Montevideo, Uruguay; Servicio Académico Universitario y Centro de Estudio Paleontológicos (SAUCE-P), Universidad de la República, Departamento de Canelones, Santa Isabel s/n, 91500, Sauce, Uruguay

**Author notes:** Corresponding author: Luciano Varela,.

**Keywords:** South America, Folivora, Cingulata, Quaternary, Macroevolution, Metabolism

## Abstract

Nutrient foramina are small openings in the periosteal surface of long bones that traverse the cortical layer and reach the medullary cavity. They are important for the delivery of nutrients and oxygen to bone tissue, and are crucial for the repair and remodeling of bones over time. The nutrient foramina in the femur’s diaphysis are related to the energetic needs of the femur, and have been shown to be related to the maximum metabolic rate (MMR) of taxa. Here, we investigate the relationship between nutrient foramen size and body mass as a proxy to the aerobic capacity of taxa in living and extinct xenarthrans, including living sloths, anteaters, and armadillos, as well as extinct xenarthrans such as glyptodonts, pampatheres, and ground sloths. Sixtynine femora were sampled, including 19 from extant taxa and 50 from extinct taxa. We obtained the blood flow index (Q_i_) based on foramina area and performed PGLS and phylogenetic ANCOVA in order to explore differences among mammalian groups. Our results show that among mammals, taxa commonly associated with lower metabolism like marsupials and living xenarthrans showed relatively smaller foramina, while the foramina of giant extinct xenarthrans like ground sloths and glyptodonts overlapped with non-xenarthran placentals. Consequently, Q_i_ estimations indicated aerobic capacities comparable to other placental giant taxa like elephants or some ungulates. Furthermore, the estimation of the MMR for fossil giant taxa showed similar results, with almost all taxa showing high values except for those for which strong semi-arboreal or fossorial habits have been described. Moreover, the results are compatible with the diets predicted for extinct taxa, which indicate a strong consumption of grass similar to ungulates and in contrast to the folivorous or insectivorous diets of extant xenarthrans. The ancestral reconstruction of the MMR values indicated a lack of a common pattern for all xenarthrans, strongly supporting the occurrence of low metabolic rates in extant forms due to their particular dietary preferences and arboreal or fossorial habits. Our results highlight the importance of considering different evidence beyond the phylogenetic position of extinct taxa, especially when extinct forms are exceptionally different from their extant relatives. Future studies evaluating the energetic needs of giant extinct xenarthrans should not assume lower metabolic rates for these extinct animals based solely on their phylogenetic position and the observations on their extant relatives.

## Introduction

Nutrient foramina are small openings in bones that allow blood vessels to enter and exit the inner parts of the bone. These foramina are clearly seen in the periosteal surface of long bones and traverse the cortical layer and ultimately reach the medullary cavity. These openings are important for the delivery of nutrients and oxygen to bone tissue, and are also involved in the repair and remodeling of bones over time (Lieberman et al. 2003; Robling et al. 2006; Eriksen 2010). Nutrient foramina can be found in almost all long bones of tetrapods, with fossil groups like dinosaurs or stem-mammals showing these openings (Seymour et al. 2012; Newham et al. 2020). The size and shape of nutrient foramina can vary widely between different species of animals, and can be influenced by a variety of factors including type and zone of long bone, body size, age, among others. In the case of the femur, previous research has shown that most bones show a reduced number of nutrient foramina (1–2) in the diaphysis of the bone, which account for 50–70% of the blood flow of the femur (Trueta 1963). The mechanical loading on the femur has a direct influence over bone formation, while both the body mass and exercise level determine the loadings suffered by the bone (Foote 1911). Furthermore, the nutrient foramina in the femur’s diaphysis are related to the energetic needs of the femur, which are largely related to the generation of microfractures due to mechanical loading stress and the consequent bone remodeling (Burr et al. 2002).

Seymour et al. (2012) showed that the area of the femur’s nutrient foramina and the femora total length can be used to calculate a relative quotient of blood flow (*Qi*) that can be compared among taxa from different taxonomic groups, allowing for a direct comparison of the energetic needs of the femur across a wide spectrum of species. Furthermore, the authors showed that the obtained estimates of blood flow, when controlled for body mass, are directly related to the aerobic capacity of taxa and, ultimately, its maximum metabolic rate (MMR). In fact, endotherms like extant mammals and birds significantly differ from ectotherms like non- varanid reptiles when comparing their nutrient foramina according to their body mass, with the former showing significantly higher intercepts in their regressions lines (Seymour et al. 2012). Considering this, several studies have used the nutrient foramina to compare the aerobic capacity of fossil taxa, as well as a proxy for the estimation of MMR in these species, including dinosaurs, birds, non-mammalian synapsids, and stem-mammals (Seymour et al. 2012; Allan et al. 2014; Newham et al. 2020; Knaus et al. 2021).

Xenarthrans are a peculiar group of mammals that are found exclusively in the Americas, with approximately 31 living species (Superina and Loughry 2015). Within the Xenarthra, sloths (Folivora) are known for their slow-moving and arboreal lifestyles, with the two extant genera (*Choloepus* and *Bradypus*) representing an exceptional case of convergent evolution (Nyakatura 2012; Delsuc et al. 2019). On the other hand, anteaters (Vermilingua) and armadillos (Cingulata) are adapted for digging and feeding on insects and other small animals, with many taxa showing considerable fossorial habits (Superina and Abba 2020). Despite their distinct morphologies and ecological roles, studies have confidently shown that all xenarthrans form a clade, and their last common ancestor was probably a myrmecophagous animal with adaptations to digging and climbing (Gaudin and Croft 2015). Furthermore, xenarthrans represent one of the four major clades within placental mammals, and their phylogenetic position has been the center of debate (Kriegs et al. 2006; Murphy et al. 2007; Morgan et al. 2013). In fact, xenarthrans could represent one of the most basal divergences within the placental mammals, making them particularly important for understanding the evolution of mammals (Svartman et al. 2006). Moreover, they show a unique set of characteristics that often represent less derived forms, but also particular adaptations product of their peculiar evolutionary history (Vizcaíno and Bargo 2014). One of these characteristics is related to their body temperature and basal metabolic rate, with studies showing members of the clade with the lowest values among mammals, oftentimes in ranges similar to those of non-placental mammals with similar habits like monotremes and marsupials (McNab 1984, 1986). In particular, sloths have received special attention due to their slow-moving and seemingly sluggish behavior. Living sloths are known for their extremely low metabolic rates and reduced muscle mass, which are thought to be adaptations to their folivorous diet and arboreal lifestyle (McNab 1985). Nevertheless, other members of the clade also present lower than expected body temperatures or basal metabolic rates, probably related to their specialized diets or fossorial habits (McNab 1985).

The fossil record of the Xenarthra is significantly more varied than the living members of the clade, both in terms of species diversity as well as morphological and ecological adaptations (Vizcaíno and Loughry 2008). In particular, within Cingulata, the glyptodonts represent a clade within the armadillo family Chlamyphoridae (Delsuc et al. 2016) that shows considerable differences with their close living relatives, being mostly giant terrestrial animals with more rigid carapaces (Fariña et al. 2013). Moreover, within sloths, several fossil forms also show tendencies towards gigantism, with several families like Megatheriidae and Mylodontidae having members with estimated body masses of more than 3,000 kg (Toledo et al. 2015). These giant ground sloths were certainly not arboreal like their extant relatives, and some of them probably had the capacity to move using only their hindlimbs (Casinos 1996; Fariña et al. 2013), form groups of individuals with a certain level of social behavior (Tomassini et al. 2020; Varela et al. 2022), or dig extensive tunnel systems (Vizcaíno et al. 2001; Frank et al. 2015). Considering the extreme differences between the fossil xenarthrans and their few extant relatives, recent research has shown that direct comparisons with the latter could not be appropriate to perform paleobiological reconstructions (Vizcaíno et al. 2018). In fact, one of the key characteristics of living xenarthrans, their low body temperature and metabolic rate, has been elusive to study in their fossil relatives since they are impossible to directly measure in commonly fossilized body parts, like bones or teeth. However, it is commonly assumed that the extinct xenarthrans, including the giant terrestrial forms like glyptodonts and ground sloths, had low metabolic rates based on their phylogenetic position (McNab 1985; Vizcaíno et al. 2023). Nevertheless, some research has suggested that some giant members of the clade could have had higher agility levels and, potentially, higher metabolic rates (Billet et al. 2013; Boscaini et al. 2018; Tambusso et al. 2021; Dantas and Santos 2022). These results could have important consequences on our understanding of the evolution of xenarthrans since they would indicate that the metabolic rates observed in most of the extant members of the clade would be the result of convergent evolution due to their particular lifestyles and not necessarily the product of an evolutionary constraint due to common ancestry.

Here, we aim to investigate the relationship between nutrient foramen size and body mass as a proxy to the aerobic capacity of taxa in living and extinct xenarthrans. Specifically, we analyzed the nutrient foramina in the femora of several species, including living sloths, anteaters, and armadillos, as well as extinct xenarthrans such as glyptodonts, pampatheres, and ground sloths. Our objectives were to determine whether the size of nutrient foramina in extant and extinct xenarthrans shows differences that could be associated to the aerobic capacity and maximum metabolic rates in these species, with particular interest in the implications on our understanding of the behavior and ecology of fossil giant xenarthrans.

## Methods

### Taxon Sampling

We extracted all the available information of mammalian taxa in the data published by Seymour (2012). Furthermore, we expanded the dataset with new data of extant mammals from specimens housed in the Statens Naturhistoriske Museum in Denmark (MHND) in Copenhagen. These new data comprised several extant xenarthrans, as well as members of other clades like marsupials and non-xenarthran placentals. Regarding the extant xenarthrans, we obtained data for the following species: the folivorans *Bradypus torquatus* and *Bradypus tridactylus*; the vermilinguans *Cyclopes didactylus*, *Myrmecophaga tridactyla* and *Tamandua tetradactyla*; and the cingulates *Cabassous unicinctus*, *Chaetophractus vellerosus*, *Dasypus hybridus*, *Dasypus novemcinctus*, *Euphractus sexcinctus*, and *Priodontes maximus*. Also, we obtained data for several fossil xenarthrans housed in the Arroyo del Vizcaíno collection (AdV), the Museo Paleontológico Profesor Armando Calcaterra (MPAC), and the Museum of Natural History (MNHN-M) in Uruguay; the Muséum National d’Histoire Naturelle (MNHN-F) in France; and the Statens Naturhistoriske Museum (MHND) in Denmark. In particular, the sampled fossil xenarthrans are represented by the following taxa: the folivorans *Catonyx cuvieri*, *Glossotherium robustum*, *Lestodon armatus*, *Megatherium americanum*, *Mylodon darwinii*, *Nothrotherium maquinense*, *Scelidotherium leptocephalum*, and *Valgipes bucklandi* and the cingulates *Dasypus punctatus*, *Glyptodon reticulatus*, *Holmesina majus*, *Neoglyptatelus* sp., *Neosclerocalyptus paskoensis*, *Neosclerocalyptus ornatus*, *Panochthus tuberculatus*, and *Propraopus sulcatus*. Sixtynine femora were sampled, including 19 from extant taxa and 50 from extinct taxa. All the newly generated data is available as supplementary material (Table S1).

### Foramen area and *Q_i_* calculation

For the analysis, we measured the nutrient foramina present in the femoral diaphysis of the studied specimens (Fig. 1). In cases where more than one foramen was present on the specimen, we measured the largest one. In the case of fossil specimens, when sediment was adhered to the bone surface and potentially interfered with the measurement of the nutrient foramen, we removed the sediment in order to completely expose its complete size. In all cases, the specimens were checked for cracks and breakages that could modify the original size of the nutrient foramen, and in those cases the foramen was not sampled. Measurements were made digitally using the software Fiji (Schindelin et al. 2012) on high-resolution digital images following Seymour (2012) and Hu et al. (2020). We measured the minor diameter of the nutrient foramen for the *Q_i_* calculation. For each specimen we also measured the femur’s total length using a digital caliper or a measuring tape for very large fossil specimens. The index of blood flow into the femur through the nutrient foramen was calculated as *Q_i_* = *r*^4^/*L*, where *r* is the nutrient foramen radius and *L* is the femur total length. Also, we obtained the average body mass (*Bm*) for each extant and extinct taxa from a literature review (Table S1).

**Figure 1.**
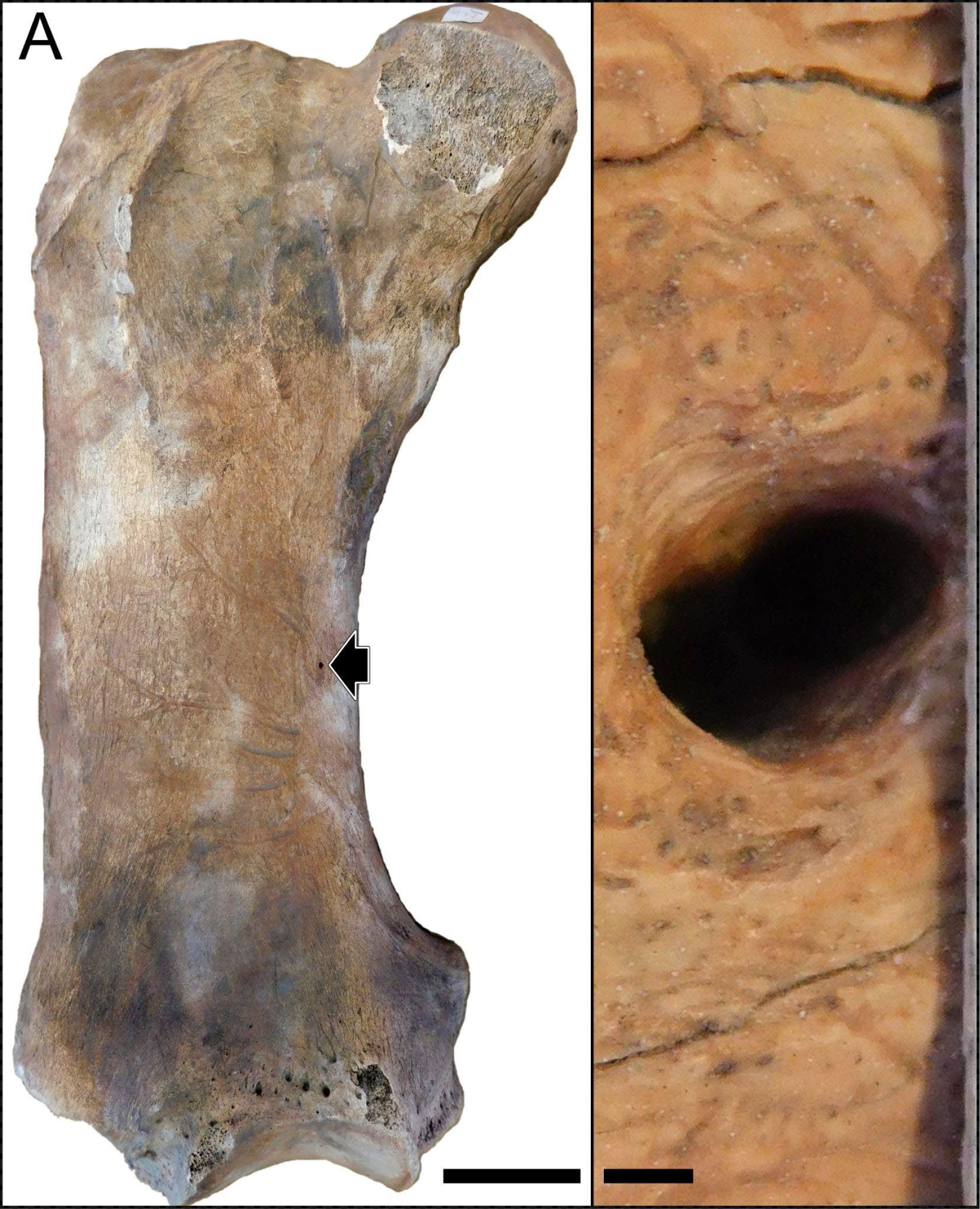
Images of the femur of a *Lestodon armatus* specimen (CAV 977). A. Entire femur in anterior view. Nutrient foramen is visible on the medial part of the diaphysis (black arrow). Scale bar equals 10 cm. B. Close view of the nutrient foramen. Scale bar equals 1 mm.

### Phylogeny

In order to account for the phylogenetic relatedness of the studied taxa in all the statistical analyses, we obtained an updated phylogeny of extant mammals from TimeTree (Kumar et al. 2022) and pruned all the taxa that were not present in our dataset. Also, we added the fossil taxa to this phylogeny following the most recent publications on fossil xenarthran systematics, namely Wible (2006); Delsuc et al. (2016), Tambusso et al. (2021) for Cingulata, Casali et al. (2020) for Vermilingua, and Varela et al. (2019), Delsuc et al. (2019), and Presslee et al. (2019) for Folivora. The final tree was constructed using Mesquite (Maddison and Maddison 2007). Since the original phylogeny was time-calibrated, the inclusion of fossil taxa resulted in a non-ultrametric tree in which the tips corresponding to older fossil taxa are consequently placed with shorter branch lengths that do not reach 0 Ma. This approach produced a phylogeny including 88 tips, with 69 of those representing extant taxa and 19 representing extinct taxa.

### *Q_i_* evolution, PGLS, and ANCOVA

The phylogenetic tree was used to address the relevance of the phylogenetic signal on the evolution of *Q_i_* and *Bm*. In particular, we fitted three evolutionary models to the data using the “fitContinuous” function of the R package Geiger (Harmon et al. 2008), namely Brownian Motion (BM), Ornstein-Uhlenbeck (OU), and White Noise (WN; i.e., absence of phylogenetic signal). We addressed model significance using Akaike weights and selected the model showing the best fit for the data for further analyses.

To establish the relationship between the *Q_i_* and *Bm* we used Phylogenetic Generalized Least Squares (PGLS) regressions, which allow to account for the non-independence of biological data due to phylogenetic relatedness in a flexible way. The flexibility of PGLS not only allows for the incorporation of several variables, but also allows for the implementation of different correlation structures like BM or OU (Harmon 2019). Considering this, we set the correlation structure according to the results of the analysis described in the previous step. These analyses were conducted using the R package “nlme” (Pinheiro et al. 2017) with the function “gls”, implementing phylogenetic weights and the correlation structure “corMartins” of the package “ape” (Paradis et al. 2004). We fitted different models accounting for the existence of differences in slopes and/or intercepts and compared them in relation to the significance of their respective coefficients, as well as using the AIC and BIC metrics. In particular, we tested the significance of a model including a categorical variable depicting a grouping based on previous assumptions about mammalian metabolism, i.e., the fact that non-placental mammals and xenarthran placentals tend to show lower metabolisms than the rest of the placentals (McNab 1984, 1986). Furthermore, in order to incorporate our hypotheses concerning fossil giant xenarthrans, we grouped both glyptodonts and ground sloths as a separate group. The normality of the residual error of the fitted models was addressed following the approach of Butler et al. (2000), by transforming the residuals by the Cholesky decomposition of the inverse of the phylogenetic covariance matrix and testing for normality of the transformed residuals using the “lille.test” function of the R package “nortest” (Gross and Ligges 2015).

In order to explore differences among the major mammalian clades, we conducted a *post hoc* test using an ANCOVA approach. Specifically, we tested for the significance of a categorical variable representing three extant mammalian groups, namely non-placentals (Prototheria and Metatheria), Xenarthra, and Epitheria (i.e., non-xenarthran placentals), and the fossil giant xenarthrans groups that are recognised as widely diverging from extant forms (i.e., Folivora, excluding both extant genera, and Glyptodontiinae). In this regard, based on the lower metabolic rates often reported for non-placentals and xenarthrans, we hypothesized that there were significant differences between their regression lines and that of the rest of the placentals. We used the “glht” function of the R package “multcomp” (Hothorn et al. 2016) and adjusted P-values according to the Bonferroni method in order to account for the effect of multiple comparisons.

### Estimation of MMR

To estimate the Maximum Metabolic Rate of the studied fossil xenarthrans, we conducted an analysis using the R package “mvMorph” (Clavel et al. 2015), which allowed us to estimate missing values for fossil tips as well as ancestral states for the mammalian phylogeny. For this analysis we fitted a multivariate OU model to the phylogeny accounting for the covariance of MMR, *Qi*, and *Bm* of taxa using the function “mvOU”. Then, we used the “estimate” function to estimate ancestral and missing tip values. In order to explore the predicting capacity of our model, we compared the estimated and observed values for taxa with MMR data and performed a linear regression to obtain a R^2^ value. This approach allowed us to obtain average and 95% confidence values for each of the studied taxa and check the reliability of the estimates against the observed values in taxa with available MMR data. We performed ancestral state reconstruction using the best model and the functions “estim” of the R package “mvMorph” and “contMap” of the package “phytools” (Revell 2012). Finally, we report Mass-independent MMR values as mLO_2_h^-1^g^-0.67^ for better comparison with previous literature (Hemmingsen 1960; Knaus et al. 2021).

## Results

### Nutrient Foramina Size and *Q_i_* in Extant and Extinct Xenarthrans

The results showed considerable variation in the nutrient foramina size across the extant and extinct xenarthrans analyzed, and, as expected, a strong correlation with body mass. Most of the studied specimens showed only one nutrient foramen in the femur diaphysis, but a minority of the specimens showed two, which were almost always located in close proximity from one another. Some fossil specimens apparently lacked any nutrient foramina in their diaphysis; however, we could not confirm if this condition was the product of sediment deposition or taphonomic alterations. Nevertheless, the absence of nutrient foramina in long bones is reported as relatively common in humans (Mysorekar 1967). The obtained *Q_i_*values for the extant xenarthrans spanned from 2.73*10^-7^ mm^3^ for the species (*Cyclopes didactylus*) to 2.09*10^-4^ mm^3^ for the largest species (*Priodontes maximus*). The *Q_i_* values for the sampled extinct xenarthrans ranged from 6.06*10^-5^ mm^3^ for the smallest armadillo (*Neoglyptatelus* sp.) to 1.72*10^-2^ mm^3^ for the largest sloth (*Megatherium americanum*). When plotting the studied extinct giant xenarthrans’ *Q_i_*against *Bm*, many taxa fell considerably above the OLS regression line of mammals, well within the expected values for placental mammals of similar size (Fig. 2).

**Figure 2.**
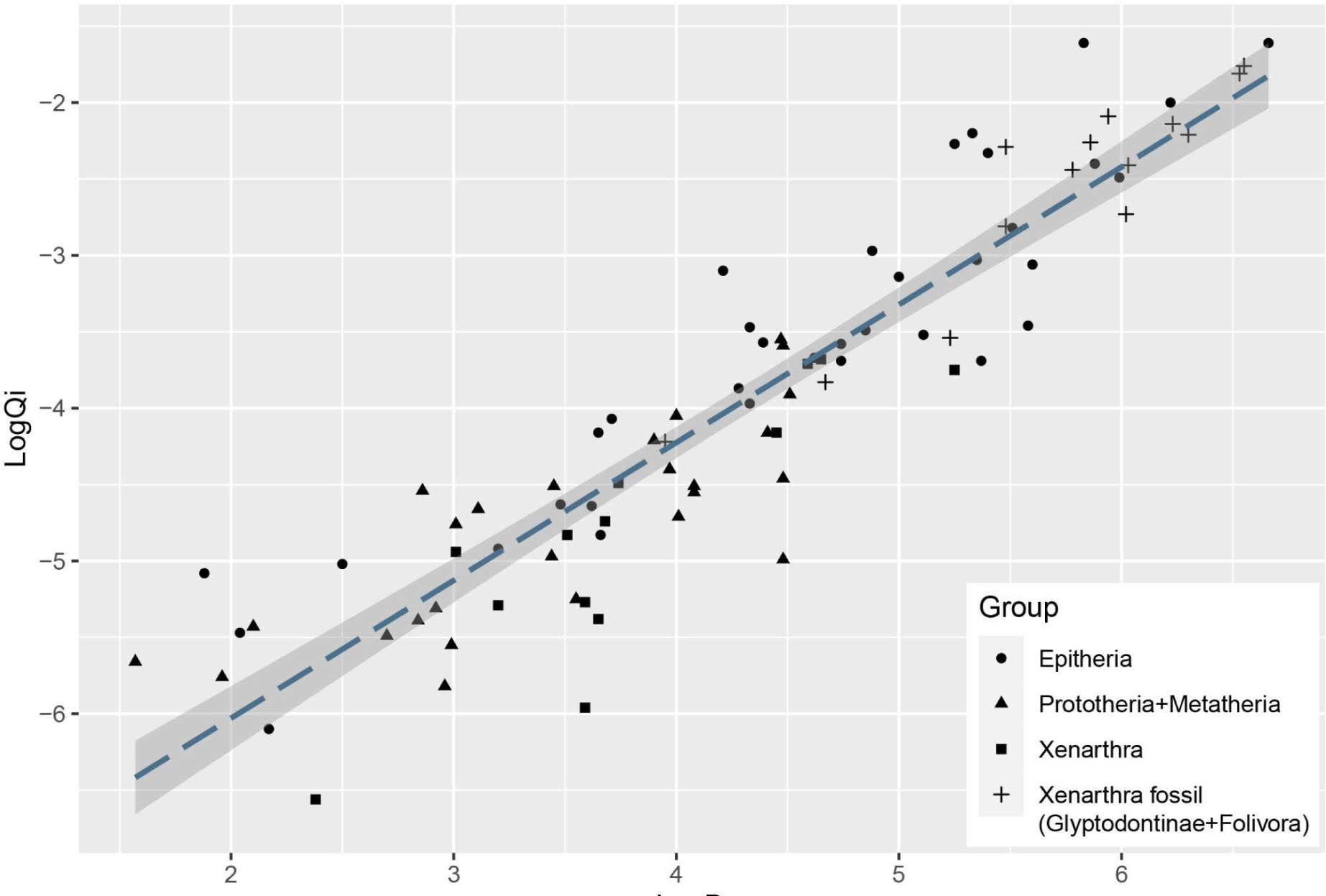
Bivariate plot of log *Q_i_* vs log *Bm* showing the placement of the different mammalian groups mentioned in the text. OLS regression is plotted for comparison with previous studies.

**Figure 3.**
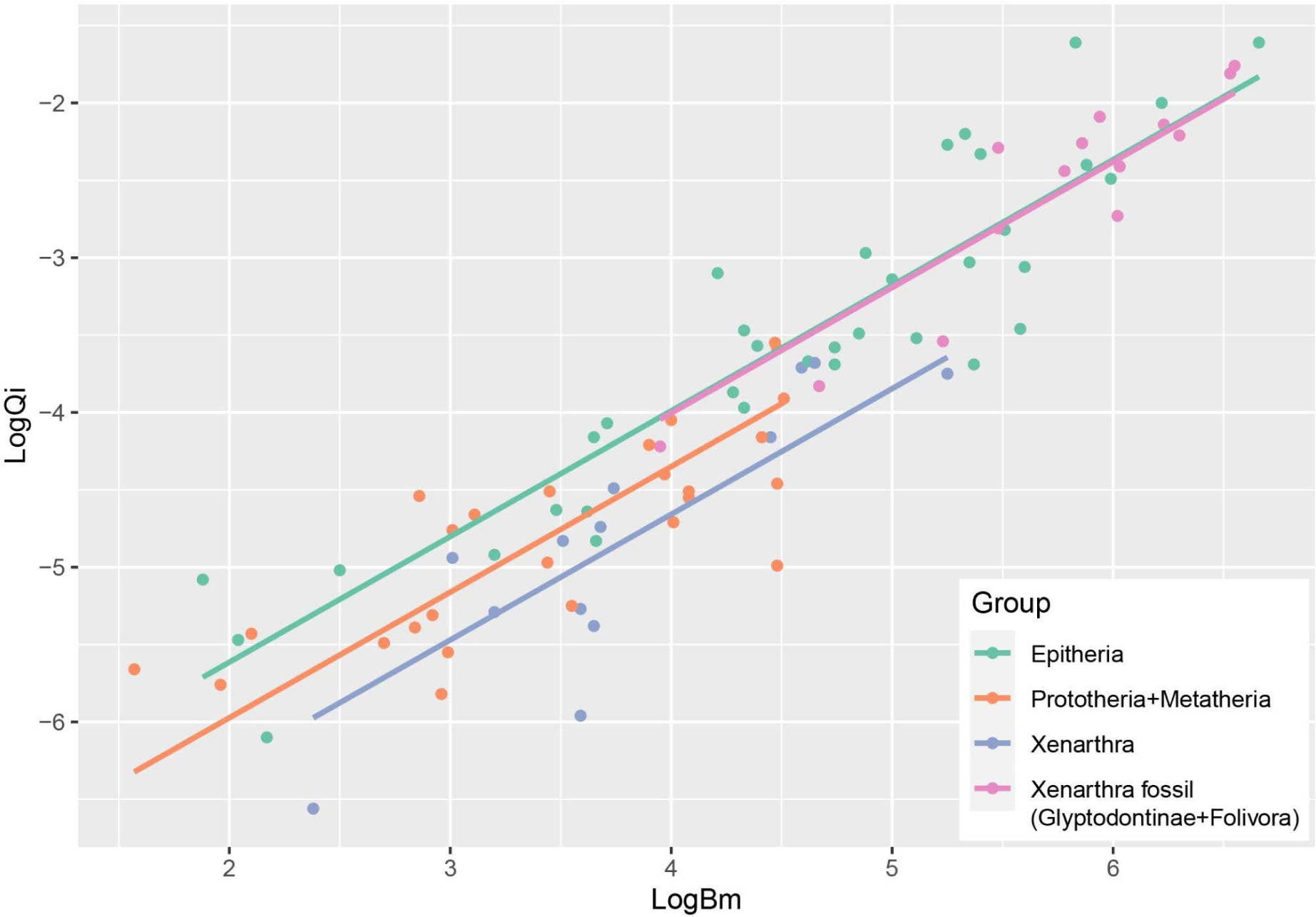
PGLS results showing the difference in intercepts recovered for the mammalian groups considered in the study.

**Figure 4.**
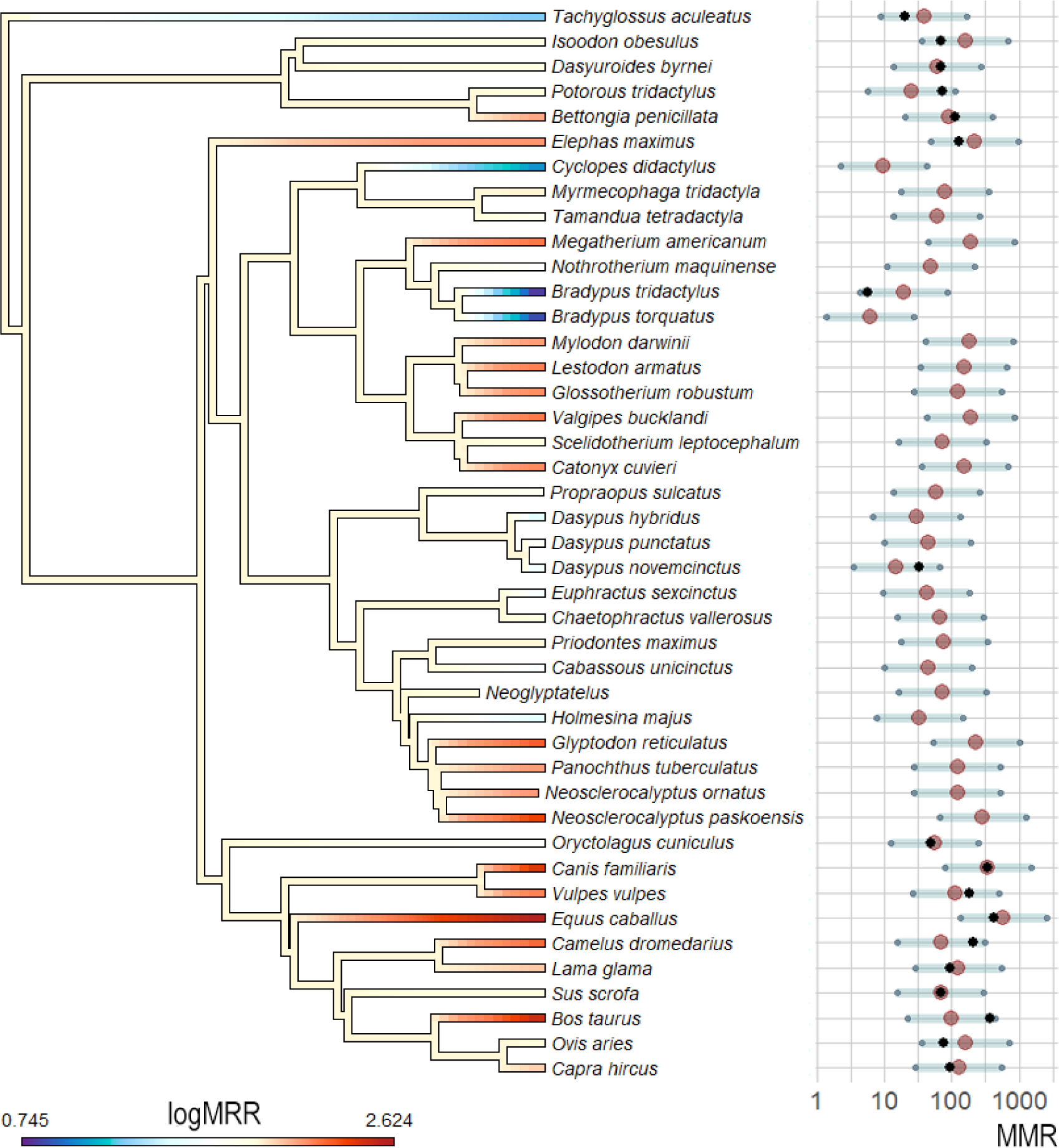
MMR phylogenetic estimations and ancestral state reconstruction using *Bm* and *Q_i_* as predictor variables and the OU model. MMR values in the phylogeny are log- transformed for better visualization. MMR average estimations and 95% confidence intervals are shown as red dots and gray bars, respectively. Black dots represent MMR values for taxa with empirical data available.

### Phylogenetic Generalized Least Squares and ANCOVA

The best fitting models for *Bm* and *Q_i_* were different. For *Bm*, the best fitting model was BM (AIC.w = 0.744), while an OU model was the best fit for *Q_i_* data (AIC.w = 0.687). In both cases, the white noise model showed extremely low fit compared to the alternatives (AIC.w < 0.001), indicating the existence of phylogenetic signals in both variables. Considering this, we used an OU model for the PGSL. As expected, the PGLS analysis showed a strong correlation between *Bm* and *Q_i_*. Furthermore, the inclusion of the categorical grouping variable in the analysis supported the distinction between some of the groups. In particular, the results showed support for the existence of different intercepts between groups (p = 0.0001), but no distinction between slopes (the interaction coefficient was not significant, p = 0.084; while the BIC showed a better fit for the model without interaction, ΔBIC > 6, and the AIC showed no preference between both models, ΔAIC < 2). The Lillefors test showed normality of the transformed residuals (p = 0.315). The post-hoc pairwise ANCOVA showed a significant difference between the Epitheria and both extant xenarthrans and non-placentals (Bonferroni-adjusted p < 0.001 and p = 0.014, respectively). On the other hand, the giant fossil xenarthrans (ground sloths and glyptodonts) showed a significant difference in intercept with the extant xenarthra (p = 0.002), but no significant difference with the Epitheria (p = 1).

### Estimation of MMR and Ancestral Character Reconstruction

The OU model was the best fitting model when compared to a BM model in the multivariate analysis (ΔAIC > 10). The observed vs predicted regression showed a R^2^ = 0.92, indicating a good prediction capacity of the model. The predicted MMR values when considering *Bm* and *Q_i_* as predictors ranged from 6.09 mL O_2_h^−1^g^−0.67^ for the sloth *Bradypus torquatus* to 589.72 mL O_2_h^−1^g^−0.67^ for the equid *Equus caballus*. The lowest predicted values corresponded to three extant xenarthrans, the sloth *Bradypus torquatus*, the anteater *Cyclopes didactylus*, and the armadillo *Dasypus hybridus*, all with values lower than 50 mL O_2_h^−1^g^−0.67^ and lower than the values registered or estimated for all the analyzed Epitheria. Epitherians showed the highest estimated values, with *Canis familiaris* and *Equus caballus* showing values higher than 300 and 500 mL O_2_h^−1^g^−0.67^, respectively. A distinction between extant xenarthrans an fossil giant xenarthrans was seen in the predicted values, with the former showing estimated MMR values lower than 100 mL O_2_h^−1^g^−0.67^ in all cases and the latter showing values higher than 100 mL O_2_h^−1^g^−0.67^ except for two sloths, *Nothrotherium maquinense* and *Scelidotherium leptocephalum*.

The ancestral reconstruction showed a pattern consistent with multiple independent cases of high or low MMRs, with almost all ancestral nodes showing intermediate MMR values. This pattern is consistent with the presence of taxa with low and high MMRs in all groups and the fact that our sample is relatively small and covers a wide spectrum of mammal species with a deep phylogenetic history. In the case of the Xenarthra, the results show an ancestor with an intermediate MMR and the independent acquisition of low MMR in the three extant clades.

## Discussion

### Size of nutrient foramina and femoral blood flow in xenarthrans

Previous research has shown consistent differences between some tetrapods regarding the size of the femur nutrient foramina, and the consequent blood flow into the femur diaphysis, and its relationship to the aerobic capacity of different groups. In particular, among extant tetrapods, birds and mammals have shown significantly larger nutrient foramina when compared to other tetrapods, which has been associated with their higher metabolic needs and more active lifestyles (Seymour et al. 2012). Furthermore, these differences have allowed researchers to study the aerobic capacity and maximum metabolic rates of fossil taxa, gaining insights regarding the metabolism of groups like dinosaurs, basal mammaliforms, among others (Seymour et al. 2012; Allan et al. 2014; Newham et al. 2020). This approach not only provides information on specific extinct taxa, but also has allowed the improvement of the understanding of the evolution of aspects such as endothermy in the fossil record (Grigg et al. 2022). In fact, it is noteworthy that some early synapsids showed evidence in favor of elevated aerobic capacity based on *Q_i_* values similar to those observed in mammals, pointing to an early evolution of the high aerobic capacity commonly seen in mammals (Knaus et al. 2021). However, no previous research has focused on the potential differences in nutrient foramina size within mammals, especially considering the large variations that exist in the clade regarding aerobic capacity and metabolic rate.

In this regard, our results show significant differences in the nutrient foramina size and the consequent inferred femora blood flow between different mammalian groups. The PGLS regressions showed considerable differences between the Epitheria and the rest of the sampled mammals, indicating that extant xenarthrans and non-placentals have proportionally smaller nutrient foramina and less blood flow in their femora compared with the Epitheria. In particular, the xenarthrans most adapted to a slow-moving lifestyle, represented by the extant sloth *Bradypus* and the vermilinguan *Cyclopes didactylus*, showed the smallest foramina sizes reported for mammals, which would be in accordance with the extremely low metabolism reported for these taxa (McNab 1978; Nagya and Montgomery 2012). On the contrary, the results showed that the extinct giant xenarthrans like glyptodonts and ground sloths had proportionately larger nutrient foramina in their femora, indicating higher aerobic capacity and agility levels when compared to their extant relatives. In fact, some of the largest xenarthrans like the sloths *Megatherium americanum* and *Lestodon armatus*, or the glyptodont *Glyptodon reticulatus* showed *Q_i_* values comparable to those observed in other giant placental mammals like elephants in our dataset.

### Aerobic capacity and maximum metabolic rate in extinct giant xenarthrans

Our results place most giant fossil xenarthra as animals with aerobic capacities just like other similarly giant placentals, such as elephants or some ungulates. In fact, two non-xenarthra taxa from the late Pleistocene megafauna, the Meridiungulata (relatively closely related to extant ungulates; Buckley 2015) *Macrauchenia patachonica* and *Toxodon platensis* also showed values similar to those of the extinct giant xenarthrans. The relative size of nutrient foramina in giant xenarthrans, as well as the estimations of MMR point to a relatively high aerobic lifestyle for these animals, which would greatly differ from that of their extant relatives. In fact, the results for *Megatherium americanum*, the largest sloth analyzed, are compatible with the previous studies mentioned before, and depict an animal probably capable of a considerable level of aerobic activity non different to what is observed in a similarly sized placental mammal like the elephant. Furthermore, most of the sampled mylodontids (the other xenarthran clade with members reaching more than 1000 kg) showed *Q_i_* and MMR values similar to those observed in placentals of similar size, which would indicate similarly aerobic capacity to *Megatherium americanum* and elephants. Interestingly, the Scelidotheriinae *Scelidotherium leptocephalum*, which is largely associated with fossorial habits based on its morphology and its association to fossil burrows (Vizcaíno et al. 2001, Patiño et al. 2021) showed lower *Q_i_* and MMR values when compared to other scelidotheriines of similar size like *Catonyx cuvieri* and *Valgipes bucklandi*. Moreover, the relatively small nothrotheriid *Nothrotherium maquinense* also showed reduced *Q_i_*and MMR values in comparison to giant terrestrial taxa. Interestingly, *N. maquinense* was a relatively small sloth interpreted as climbers that probably fed on leaves, which would be compatible with our results based on what is known from extant sloths (Dantas and Santos 2022; Santos et al. 2023). Regarding cingulates, it is interesting to note that fossil armadillos closely related to extant taxa, like *Propraopus sulcatus* and *Dasypus punctatus*, and morphologically similar pampatheres, like *Holmesina majus*, showed overlapping MMR values with extant taxa, even in cases where extinct forms are considerable larger that extant ones. These fossil taxa are often depicted as capable burrowers and probably had similar diets and habits to those observed in extant armadillos, which are characteristics related to the low metabolic rates observed in extant xenarthrans. On the other hand, giant glyptodonts showed elevated Q_i_ and MMR values compatible with higher aerobic capacities and metabolic needs. These findings are interesting considering glyptodonts are often depicted as slow-moving, heavy-weighted, and armored taxa that probably were not capable of agile movements (Gillette and Ray 1981). However, the results would be more in line with predictions based on certain morphological traits seen in glyptodonts that proposed the existence of intraspecific fights using their club-like tails (Alexander et al. 1999). In fact, one of the glyptodonts that showed high MMR values was *Glyptodon reticulatus*, which has been proposed as capable of acquiring a bipedal stance in agonistic contexts (Fariña 1995).

Most giant fossil xenarthrans probably presented extreme departures from the common habits and morphology observed in their closely-related extant forms. In particular, some fossil taxa show extreme gigantism, which impedes the consideration of extant forms as potential analogs for their study (Vizcaíno et al. 2018). Furthermore, their large body size would also have had a significant impact over other aspects of these taxa, including their behavior, ecology, and evolution. Moreover, aspects like the extreme modification of the carapace in giant glyptodonts or the inability to climb trees of giant ground sloths further increases their unlikeness to extant forms. On the other hand, these fossil xenarthrans have been associated with what is known of their extant relatives, often depicting them as slow and not agile (Gillette and Ray 1981; Toledo 1996). Likewise, low body temperatures and metabolic rates have been assumed for extinct giant xenarthrans mostly based on their phylogenetic position (McNab 1985).

However, some research has shown evidence in favor of a more active lifestyle for some extinct xenarthra in comparison to their extant relatives. For example, several studies concerning the intracranial morphology of extinct xenarthrans, in particular the auditory region, have shown evidence in favor of a more active lifestyle in comparison with extant members of the clade. Billet et al. (2013) studied the inner ear of several xenarthrans and showed that the megatheriid *Megatherium americanum* was more agile than extant sloths based on the scaling of its semicircular canals. Boscaini et al. (2018) recovered similar results for the mylodontid *Glossotherium robustum*, arguing that this ground sloth had an inner ear morphology more similar to terrestrial taxa like *Tamandua* than to other xenarthran with fossorial habits. In fact, Boscaini et al. (2018) results placed *G. robustum* among mammals of similar size categorized as having a “medium” level of agility according to Spoor et al. (2007). Furthermore, another study by Tambusso et al. (2021) noted that extinct glyptodonts like *Glyptodon reticulatus*, *Panochthus tuberculatus*, and *Doedicurus clavicaudatus* would fall within the agility range observed in most extant Cingulata and Vermilingua, but considerable overlap among agility categories was present based on semicircular canals’ morphology. These studies would indicate that giant fossil xenarthrans could have had aerobic capacities comparable to other giant mammals in opposition to most of the extant members of the clade like sloths or fossorial armadillos. Other studies also point in a similar direction, indicating the possibility of considerable aerobic capacity. For example, the crural index of the fossil sloth *Pyramiodontherium scillatoyanei* was considerable high when compared to other xenarthra, with a value similar to those found in ungulates like llamas and horses, potentially indicating a relatively high agility (De Iuliis et al. 2004). Furthermore, the estimation of *Megatherium americanum* walking speed based on fossil tracks by Casinos (1996) and Blanco and Czerwonogora (2003) showed a range of 0.8 to 2.2 m/s, which would be similar to the normal walking speed reported for Asian elephants of similar size (1.37 ± 0.28 ms^−1^; Ren et al. 2008). For glyptodonts, some studies have proposed the existence of interspecific fights, as well as a certain capacity to adopt bipedal locomotion (Fariña et al.1995; Alexander et al. 1999; Vizcaíno et al. 2011).

### Implications for the understanding of fossil giant xenarthrans metabolism

Regarding the metabolic rate of fossil xenarthrans, most studies have focused on exploring the basal metabolic rate of members of the group. However, minimum and maximum metabolic rates are correlated in vertebrates and, therefore, our results provide important insights that can be compared to previous research (Auer et al. 2017). A study by Vizcaíno et al. (2006) showed some differences in the occlusal surface areas of fossil giant sloths, with *Megatherium americanum* having higher than expected oral processing capacity, pointing to high energetic needs in this taxon, while mylodontids showed lower processing capacities. A recent study by Dantas and Santos (2022) expanded this approach and showed results consistent with high metabolic rates for most of the Brazilian Intertropical Region fossil ground sloths. In particular, the results showed probable high metabolic rates for members of the Mylodontidae, Megatheriidae and Megalonychidae, while the Nothrotheriidae showed lower values comparable to those of extant xenarthrans. Other studies have also pointed to probable high metabolic rates in fossil sloths based on different methods. Recently, Tejada-Lara et al. (2021) proposed that the interpretation of low metabolic needs in fossil mylodontids could be instead explained by the consumption of higher-quality food like meat based on the finding that *Mylodon darwinii* probably consumed animal protein according to Isotope data from amino acids, in line with other proposals of animal items in the diet of Pleistocene sloths (Fariña 1996; Fariña and Blanco 1996; Fariña and Varela 2018). Furthermore, this interpretation would also explain the capacity of some sloths like *M. darwinii* to have lived in fairly extreme cold weathers like southern Patagonia (Varela and Fariña 2016; Varela et al. 2018). Similarly, the megalonychid *Megalonyx jeffersonii* has been recovered in northern North America, with a latitude record at 68°N in Canada, indicating that this species could also endure considerable cold climates (Harington 1978). Other indirect evidence that would point to higher metabolic rates in fossil sloths in comparison to extant forms could be related to the evolution of a complete marine lifestyle in the nothrotheriid genera *Thalassocnus* (Amson et al. 2014). In this aspect, many extant marine mammals like odontocetes, otariids, and sea otters show elevated metabolic rates for their body mass when compared with terrestrial forms (with the clear exception of sirenians, which show lower metabolic rates for their body mass; Costa 2009). Furthermore, besides their arboreal and fossorial habits, the low metabolism seen in extant xenarthrans has also been related to their particular diets. McNab (1985, 1986) showed that the constraints imposed by the diets of extant xenarthrans (myrmecophagy and folivory) were important determinants for their low body temperatures and metabolic rates. Concerning this, it is important to note that many, if not all, fossil giant xenarthrans were probably adapted to other food sources. For example, many studies have shown that most of the late Pleistocene mylodontids were adapted to the consumption of grass, with many taxa probably consuming high percentage of grass similar to ungulate grazers (Bargo and Vizcaíno 2008; Varela et al. 2023a; Varela et al. 2023b). In fact, these adaptations would be favored by a developmental mechanism in mylodontids, where the posterior teeth would be enlarged due to an unique inhibitory cascade pattern within sloths (Varela et al. 2020). In this scenario, the mere assumption of low metabolic rates for extinct giant xenarthrans based solely on their phylogenetic relatedness to extant forms is not supported, and studies should contemplate metabolic rates equal to those of other Eutheria when studying giant fossil xenarthrans.

Finally, based on the ancestral reconstruction of MMR, it would seem that the low metabolism often reported for extant xenarthrans would not represent a trait acquired early in the evolution of the clade, but, more probably, a later and independent acquisition. Interestingly, these low metabolic rates could be the product of the ecological preferences of the extant xenarthrans, representing an exemplary case of convergent evolution due to peculiar habits shared by almost all extant members of the clade, like insectivory (and myrmecophagy in some cases) and folivory, or by contrasting habits like the fossoriality of armadillos or the suspensory habits of sloths, which are both associated to lower metabolic rates (McNab 1985). Considering this, extant xenarthrans would not represent good analogues for studying the metabolism of extinct taxa. Likewise, the fact that extant xenarthrans show relatively low agility levels and metabolic rates should not be considered as evidence for the presence of these characteristics in extinct taxa, especially in giant forms like glyptodonts and ground sloths. This has important implications when addressing different ecological questions like the roles these giant animals had in past ecosystems (Vizcaíno et al. 2023). For example, Fariña (1996) made his calculations about the thermodynamics of the trophic relationships in Lujanian mammals using values of metabolic rates according only to body size, i.e., without phylogenetic correction or any other variable, which would be reasonable for the giant xenarthrans of this fauna considering our results. Furthermore, more recent and refined calculation did use those corrections without major changes in the results (Fariña et al. 2013).

## Conclusions

The study of physiological traits like metabolism is often difficult in fossil taxa due to the common lack of preservation of soft body parts in the fossil record. Considering this, it is indispensable to explore morphological features that can be associated to species’ metabolic rates but also commonly present in the fossil record. In this regard, the study of femora nutrient foramina represents an excellent opportunity to explore the aerobic capacity and maximum metabolic rate in extinct species without clear extant analogues like the giant xenarthrans. Despite this, the results of these kinds of analyses have some limitations and the limited data on extant taxa’s MMR reduce models capacity to estimate extinct taxa’s values. Future research on extinct giant xenarthrans should focus on expanding the available MMR data for extant taxa, but also on incorporating other variables associated with metabolic rate in fossil taxa to further improve our understanding of these strange animals.

## Declaration of competing interest

The authors declare that they have no known competing financial interests or personal relationships that could have appeared to influence the work reported in this paper.

## Data availability statement

All the data used in this paper is available either in the main text or the supplementary materials. The code used in the study is provided in: https://github.com/lvar/Varela_et_al._2023

## Authors contribution

**Luciano Varela**: Conceptualization, Investigation, Methodology, Formal analysis, Resources, Visualization, Writing – Original Draft. **P. Sebastián Tambusso**: Resources, Writing – Review & Editing. **Richard A. Fariña**: Resources, Funding acquisition, Writing – Review & Editing.

## Acknowledgments

We thank the curators of the following collections, who kindly allowed us access to the specimens under their care: Kasper Lykke Hansen (NHMD); Guillaume Billet (MNHN.P), Andrés Rinderknecht (MNHN-M); and the personnel of the museums of Colonia del Sacramento, Uruguay. This work was supported by a grant from the Agencia Nacional de Investigación e Innovación (ANII; FCE_1_2019_1_156563).

## Notes

### Competing Interest Statement

The authors have declared no competing interest.

